# Trends and drivers of prairie stream fish community persistence: implications for climate refugia

**DOI:** 10.64898/2026.09.08.750286

**Authors:** Niall G. Clancy, Frank J. Rahel, Annika W. Walters

## Abstract

Identifying the characteristics of climate refugia for multiple species is becoming increasingly urgent. We used repeated, fish community surveys in streams of the northern Great Plains, USA to characterize occupancy change between historical and contemporary surveys. We then identified variables correlated with community change. Across all sites, average community turnover was 56%, native species persistence was 60%, and contemporary surveys consisted of 37% colonizing species (native or introduced). Community turnover was driven about equally by species loss and colonization. Individual species showed highly variable trends in occupancy with native plains minnow (*Hybognathus placitus*), northern redbelly dace (*Chrosomus eos*), longnose sucker (*Catostomus catostomus*), and flathead chub (*Platygobio gracilis*), declining substantially, and introduced smallmouth bass (*Micropterus dolomieu*) and native channel catfish (*Ictalurus punctatus*) increased considerably. While factors associated with individual species persistence and colonization varied, warm temperatures and agriculture were most associated with community change. Importantly, cool water temperatures appear to buffer fish communities from turnover at sites moderately impacted by agriculture. Protection of cool-water refugia will thus be critical for persistence of fishes in agricultural regions.

## INTRODUCTION

A common fisheries management objective is persistence of populations and communities through time. However, warming temperatures, invasive species, human development, stream fragmentation, and drying are threatening freshwater fish communities (Perkin et al. 2015; Comte et al. 2021a; Rumschlag et al. 2025). These threats are disrupting historical environmental conditions, and communities are shifting in response (Comte et al. 2021a; Rumschlag et al. 2025). Communities that persist for long-periods of time despite these changes may reflect the conditions necessary for climate refugia.

Identifying and conserving climate refugia for vulnerable species is required to stem the biodiversity crisis (Morelli et al. 2020). Climate refugia are habitats with conditions that remain suitable for vulnerable species over long time periods, buffering them from the effects of climate change (Keppel et al. 2024). Accurate delineation of refugia is complicated by unknown species requirements, species interactions, and habitat fragmentation (Barrows et al. 2020). This context dependency makes predicting the locations of refugia difficult.

There is ongoing debate about the degree to which fisheries management should prioritize conservation of persistent coldwater refugia (Isaak et al. 2015; Isaak and Young 2023) to the neglect of connectivity that promotes optimal growth and stable metapopulations (Armstrong et al. 2019; Hahlbeck et al. 2020). Movement of fishes between seasonally suitable habitats is certainly critical for community resilience, especially in non-perennial riverscapes (Falke et al. 2011; Marshall et al. 2016). However, invasive species can make otherwise suitable refugia untenable for long-term persistence (Alofs et al. 2014), so increasing habitat connectivity may come at the cost of allowing invasive species to spread (Rahel et al. 2008; Kirk et al. 2022). Further, the influence of different land-uses is largely absent from these conversations, likely because climate refugia studies have primarily focused on salmonid populations in relatively undisturbed landscapes. No study has examined all these variables at once to help managers determine whether to prioritize connectivity or local habitat management for fish community persistence and conservation of climate refugia. As illustrated by the potential trade-offs between connectivity and invasive species, considering multiple factors that may be important for climate refugia and their interactions is critical for promoting population and community persistence.

Fish communities in the Great Plains are well suited for addressing questions about climate refugia and community persistence because they contain a diverse array of native and introduced species and exist along strong gradients of natural and anthropogenic disturbance. However, prairie systems, especially at northern latitudes, receive far less research attention than streams in other regions (Lenhart et al. 2023). Cattle grazing is the dominant land-use in the northern and western Great Plains with cropland agriculture being significant in some areas. Major threats to stream fishes are groundwater withdrawal, habitat alteration, stream fragmentation, and introduced species (Bramblett et al. 2005; Perkin et al. 2015; Coulter et al. 2024). Stream warming and drying are also growing threats to prairie fishes (Kirk and Rahel 2022; Clancy et al. 2025; Rieger et al. 2026).

We analyzed a multi-decade dataset of 279 repeated fish-community survey sites across the northern and central Great Plains of the United States (Figure 1) to identify trends in site occupancy and determine what combination of local habitat, biotic characteristics, and stream connectivity corresponds with community persistence and likely climate refugia. Our objectives were to (1) quantify net changes in occupancy for individual species and community composition; (2) identify environmental characteristics of sites associated with individual species’ persistence or colonization; (3) identify environmental characteristics of sites with low rates of community turnover, high overall native-fish persistence, and low rates of colonization by both native and introduced species; and (4) determine if community change metrics and their environmental drivers varied across the study area.

**Figure 1.**
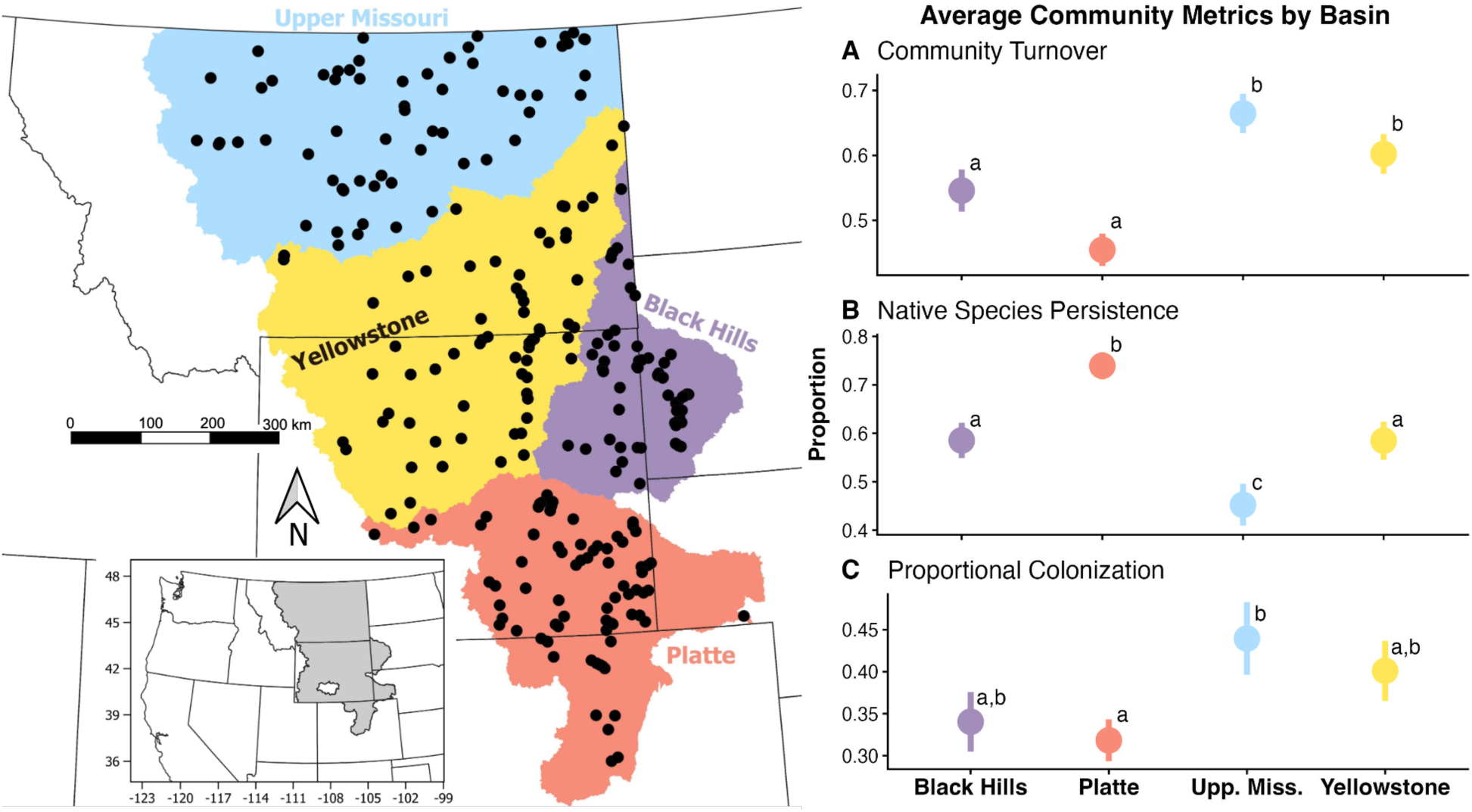
(Left) Map of repeated survey sites (black dots) in the study area with major basins differentiated by color. The Cheyenne and Little Missouri River basins were combined to make the “Black Hills” basin. (Right) Mean community metrics for each major basin. Proportion refers to the average proportion across sites for the metric titled above each graph: A) community turnover (proportion of species that were extirpated or were colonists), B) native species persistence (proportion of native species persisting), and C) proportional colonization (proportion of species colonizing). Error bars represent standard errors. Lowercase letters next to points represent basin metrics that were not significantly different from one another.

## METHODS

### Study Area

Our study area was a large region of the west-central United States spanning the northwestern portion of the Missouri River basin (Montana, Wyoming, North Dakota, South Dakota, and Nebraska) – a region containing the upper Missouri River, Yellowstone River, greater Black Hills (Little Missouri and Cheyenne River basins), and Platte River basins (Figure 1). This diverse area is home to wide expanses of prairie and sagebrush-steppe rangelands with island mountain ranges interspersed throughout.

### Fish Survey Data

We combined multiple fish-occurrence datasets (Clancy et al. 1980; Propst 1982; Barfoot 1993; EPA 2016, 2023; Comte 2021b; FishMT 2024; Haworth and Bestgen 2024; Clancy 2024; Rieger and Clancy 2025) and extracted all survey events in which full fish communities were sampled via the following criteria: (1) at least three fish species were captured, one of which was a small-bodied, nongame fish, or (2) the sample event was from a study that specifically stated all captured fishes were recorded. We then split this community-level dataset into a “historical” subset (sampled from 1934-2004) and “contemporary” subset (sampled from 2004-2024). While arbitrary, we chose 2004 as a cutoff so that there would be at least 20 years of contemporary surveys. We limited the dataset to locations sampled in both time periods, considering a site resampled if it was within two linear kilometers of the historical location on the same stream, was sampled at least nine years after the first historical sample, and used a similar type of collection gear, if recorded (historical gear recorded at 56% of sites). At sites where multiple surveys were available during either the historical or contemporary subsets, we compared the earliest possible survey to the latest possible survey. We further limited surveys to those from streams with mean August streamflow below 2.8 m^3^/s (100 cfs) due to the high likelihood of gear bias in large streams. This resulted in a preliminary dataset of 225 resampled locations. To spatially balance our dataset, we surveyed an additional 54 historical sampling locations in 2024, replicating the gear and effort used as best as possible and also attempting to sample within two kilometers of the historical location (Figure 1). A field notebook and voucher specimens were deposited at the University of Wyoming Museum of Vertebrates (https://arctos.database.museum/project/10004957). The final dataset had 279 resurveyed sites with a mean of 26 years between surveys (median = 23, standard deviation = 14, range = 9 - 86). For each species found at a site (either historically or contemporarily), a presence-absence trend value was assigned corresponding to extirpated (-1; historically present, contemporarily absent), persisted (0; historically and contemporarily present), or colonized (1; historically absent, contemporarily present). This work was approved by the University of Wyoming Institutional Animal Care and Use Committee (#2022-0102).

### Species and Community-Level Changes Over Time

#### Quantifying individual species change

From our dataset of 279 repeated fish survey sites, we calculated a mean occupancy trend (between historical and contemporary surveys) for each individual species present at a minimum of 20 unique sites as the mean of all site-level presence-absence values for that species (-1, 0, or 1) (Table 1). As such, a mean occupancy trend equal to 0 indicates no change between time periods (the species remains at the same number of sites). Calculations for native species were limited to sites within their native range.

**Table 1.**
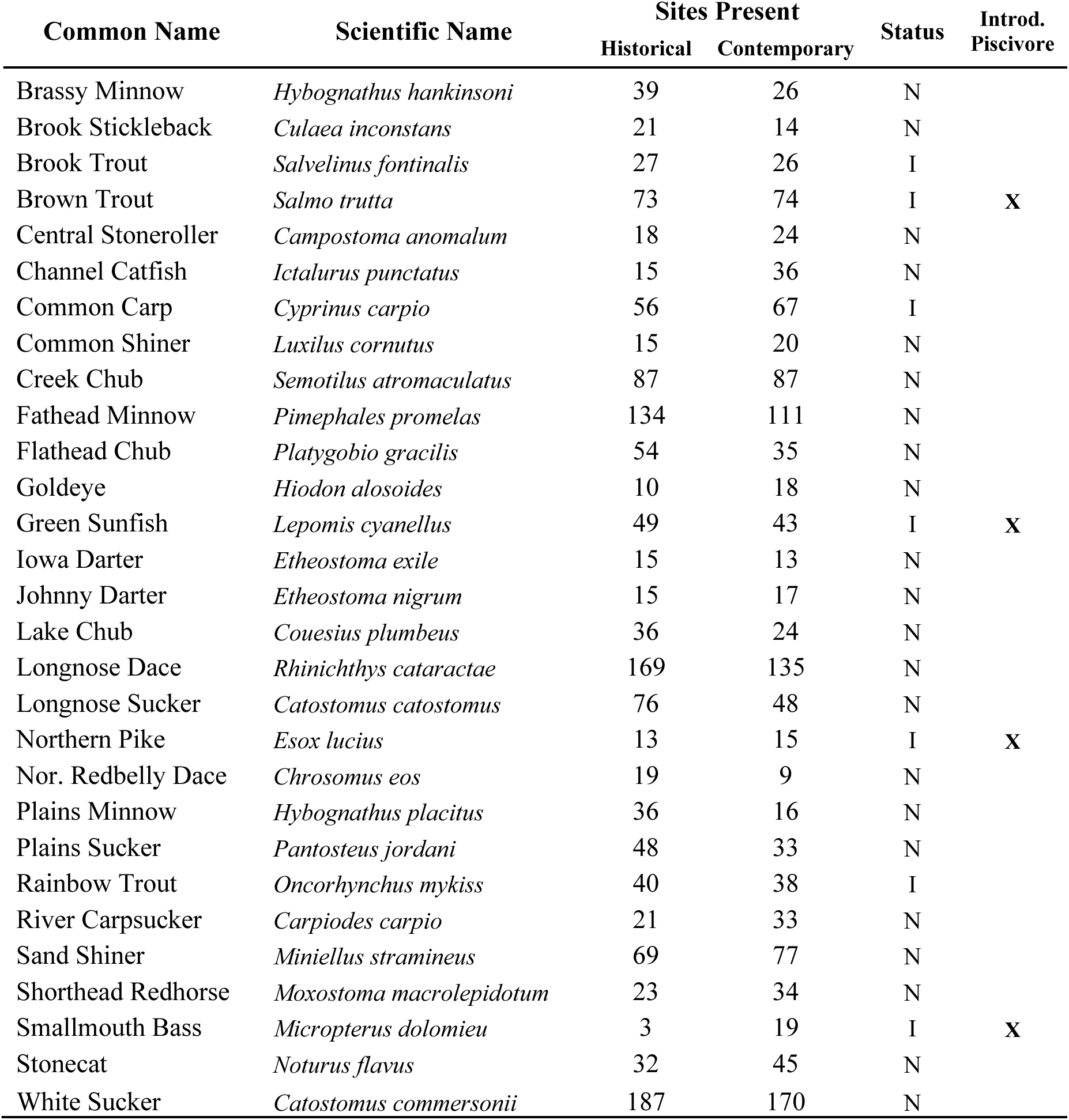
Species common and scientific names, number of sites present in the historical and contemporary time periods, native status, and whether or not the species is an introduced piscivore. Only species found at a minimum of 20 unique sites are shown. Status N indicates the species is native to some (or all) of the study area and I indicates the species was not native anywhere in the study area. The last column highlights introduced species that are piscivorous as adults.

We also calculated a response metric for use in regression analyses (see below) for individual species persistence/colonization (sites where a species was found during the contemporary survey regardless of presence or absence in the historical survey). Because this metric was intended as the response variable for multivariate regressions, it was only calculated for species present during the contemporary survey at a minimum of 30 sites to minimize model overfitting.

#### Quantifying community change

Community change metrics were those we thought would best inform refugia delineation: (A) mean community turnover, (C) mean native species persistence, and (D) mean proportional colonization.

All species, regardless of number of sites present, were included in community-level calculations. Community turnover (i.e., Jaccard’s dissimilarity, Jaccard 1912) is the proportion of species in a community that were either extirpated or colonized between two time periods. Mean community turnover was thus calculated as:

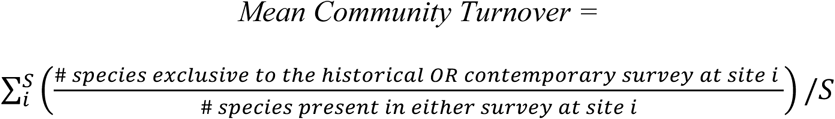

where S is the total number of sites. Values range from 0 (no turnover, i.e., same species present in both time periods) to 1 (complete turnover, i.e., no species in common between time periods).

Native species persistence is the number of native species present at a site during both the contemporary and historical surveys divided by the number of those species present only during the historical survey of the site. We then calculated Mean Persistence as the proportion of native species that remained present at a site averaged over all sites:

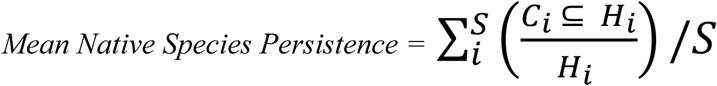

where *H_i_* is the number of native species present during the historical survey of site *i, C_i_* is the subset (<u>⸦</u>) of *H_i_* found at the site during the contemporary survey, and S is the number of sites with at least one native species present during the historical survey. Values can range from 0 (no native species persisted) to 1 (all native species persisted).

Proportional colonization was calculated as the number of species new to a site during the contemporary survey divided by the total number of species present in the contemporary survey. Mean proportional colonization was thus calculated as:

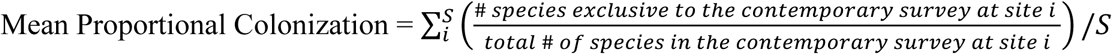

where S is the number of sites with at least one species present in the contemporary survey. Value of 0 means no new species colonized the site, and 1 means that all of the species at the site were colonists.

### Drivers of Species- and Community-Level Change

#### Predictor Variables

We obtained seven predictor variables for each resurveyed site that we thought may influence fish communities: mean August stream temperature (°C; 1993-2011 average from NorWeST, Isaak et al. 2017); stream size (m^3^/s; mean August discharge, US Forest Service Streamflow Metrics dataset; Wenger et al. 2010; USFS 2022); flow permanence (mean probability of flow presence from 1989-2021, USGS PROSPER model; Sando et al. 2022); stream fragment length (stream kilometers between barriers on the same stream or the stream mouth (where the stream flows into different stream), derived from the national hydrography dataset using QGIS); presence (1) or absence (0) of a downstream dam or perched culvert on the same stream (SARP 2024 barrier inventory); introduced piscivore presence (1) or absence (0) during the contemporary (but not necessarily historical) survey (Table 1); and areal proportion of cropland or pastureland within a 5 km radius of the survey site (derived using QGIS from the 2024 National Landcover; USGS 2024). Introduced piscivores were classified based on having substantial evidence of piscivory amongst stream-dwelling populations. The most commonly captured were brown trout (Budy and Gaeta 2017), green sunfish (Lohr and Fausch 1996), northern pike (Hogberg 2024), and smallmouth bass (Gard 2004). We examined aerial imagery of each purported dam or perched culvert and removed those that were clearly not fish barriers (e.g., due to the presence of major side channels or no obvious barrier being present). Barriers were not included if they formed a substantial reservoir because reservoirs likely serve as a colonization source for many species. Flow permanence estimates were not available for 11 sites, so we imputed values for nine sites based on a linear regression from all sites having PROSPER values using elevation and stream size as predictors (r^2^ = 0.111, p < 0.01). The two remaining sites were given the same value as sites within five km on the same stream. Multicollinearity of predictor variables was assessed, and none were substantially correlated (|r| < 0.3).

#### Regression Analyses

To determine which predictor variables correlated with response variables, we used binomial logistic regression models implemented in program R (R Core Team 2024) with the assistance of ChatGPT version 5.1 (OpenAI 2025). Initial models that included the number of years between samples as a predictor showed no significant relationship with the response variables (lowest p-value = 0.47), so number of years was not included in further models. The full-variable model for each logistic regression was thus

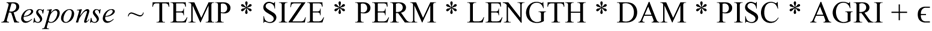

where TEMP is mean August stream temperature, SIZE is mean August stream discharge, PERM is the probability of flow permanence, LENGTH is length of the stream fragment, DAM is the presence or absence of a dam downstream of the survey site (on the same stream), and PISC is the presence or absence of an introduced piscivore (Table 1), AGRI is the proportion of land within 5 km of the survey site classified as cropland or pastureland (referred to as agricultural land use), * indicates full interactions between variables, and ɛ is the model error structure. Models for community turnover and proportional colonization did not include introduced piscivore presence because those piscivores were included in the calculation of those metrics.

Best-fit models (including an intercept only model) for each community response were identified using AIC. Models without interactions were also run, and the seven individual predictors were scaled so we could determine individual variable importance. We constructed variable importance plots of the full-variable model for each response.

#### Species Contributions to Mean Community Turnover

To determine whether species loss or species colonization was primarily responsible for community change and which species were driving change, we also calculated two metrics of proportional contribution to mean community turnover. First, the mean proportion of turnover due to colonization was calculated:

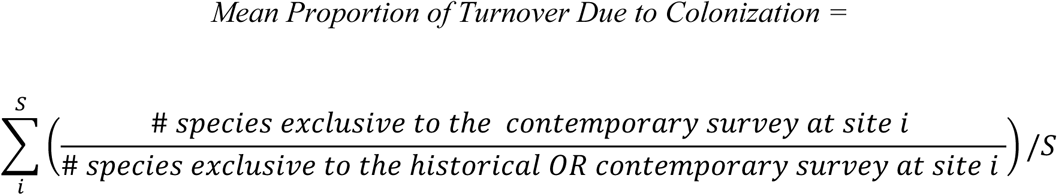

where *S* is the total number of sites. The mean proportion of turnover due to species loss is therefore: 1 – (the mean proportion of turnover due to colonization).

Second, the mean proportion of turnover due to each individual species was calculated:

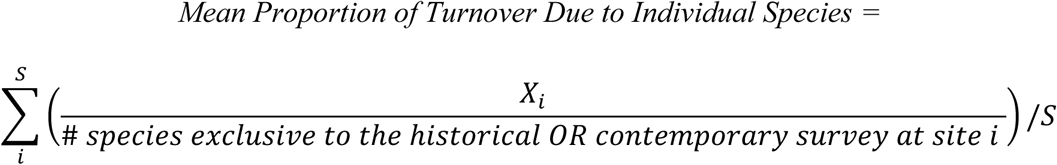

where *X* = 1 if the species colonized or was lost at site *i*, and X = 0 if the species persisted or was not present at site *i*.

### Drivers of Community Change Across Basins

Variable importance was also determined separately for the three community response metrics within each major basin (Figure 1). As before, variable importance was determined from logistic regressions without interactions, and models for community turnover and proportional colonization did not include piscivore presence. Mean community-metric differences between the basins were assessed with an analysis of variance (ANOVA) and a Tukey test.

## RESULTS

### Individual Species Occupancy Between Time Periods

Individual species showed high variation in mean occupancy trends between time periods (Figure 2). Eleven species were declining at a statistically significant level (standard error bars not overlapping zero). All 11 were native species. Eight species were increasing at a statistically significant level, six native, two introduced. Trends for the remaining 10 species were not significantly different from zero.

**Figure 2.**
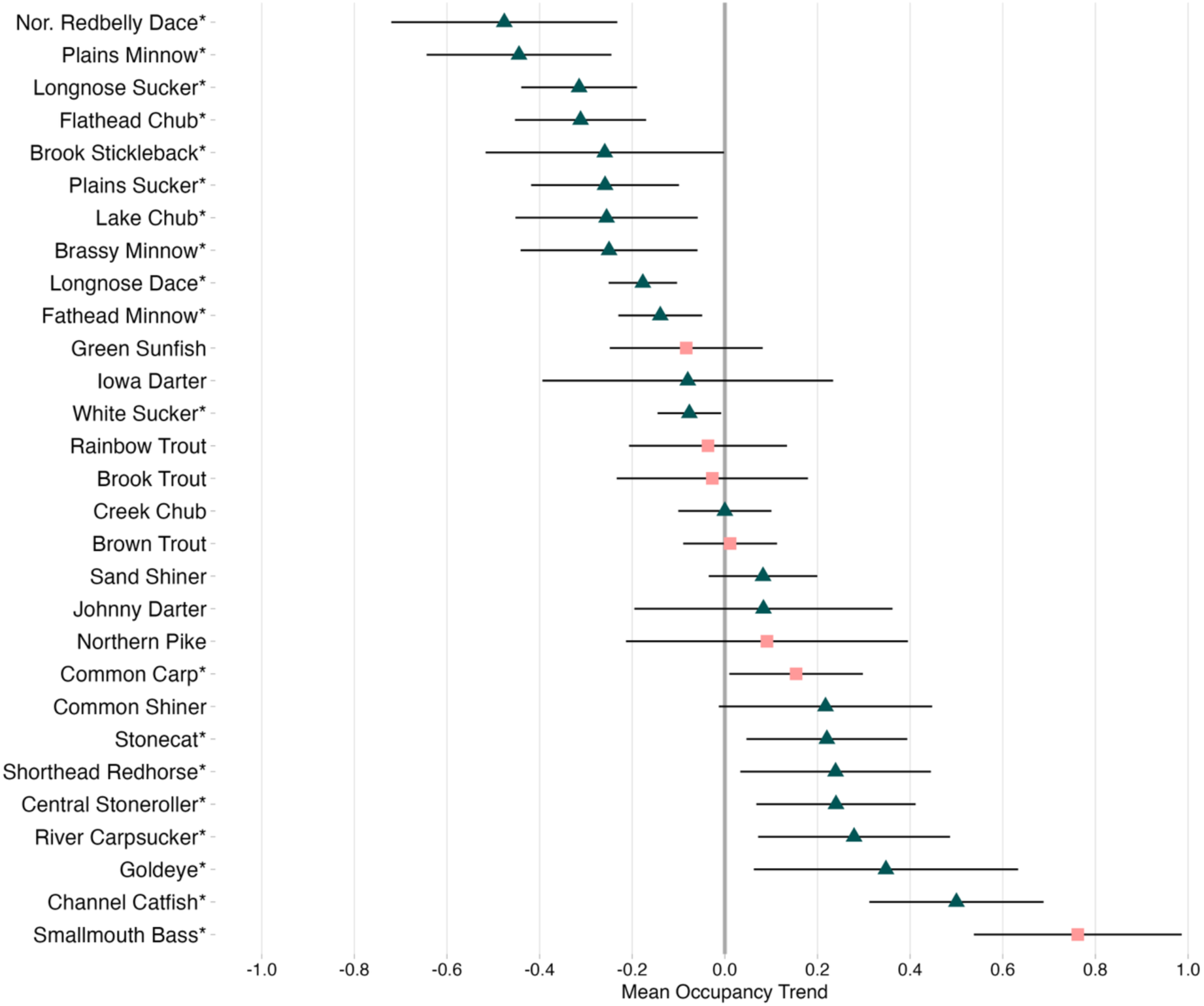
Species’ mean occupancy trends between historical and contemporary time periods. The dark grey line indicates no net change in number of sites occupied. Error bars represent 90% confidence intervals where only those not overlapping 0 (dark grey line) are considered significantly increasing or decreasing and are denoted by an asterisk. Only species found at a minimum of 20 unique sites are shown. For native species (green triangles), analysis was limited to the native range only.

### Drivers of Species and Community-Level Change

The most important variables associated with species persistence and colonization differed among species, but temperature was the most common, significant predictor (9 of 20 species) followed by agricultural land-use and fragment length (both 4 of 20 species) (Table 2). Top predictors (whether or not they were statistically significant) for individual species persistence and colonization were as follows: temperature for seven of 20 species, agricultural land-use for four species, downstream barrier presence and fragment length for three species each, stream size for two species, piscivores for one species, and streamflow permanence for none (Table 2).

**Table 2.**
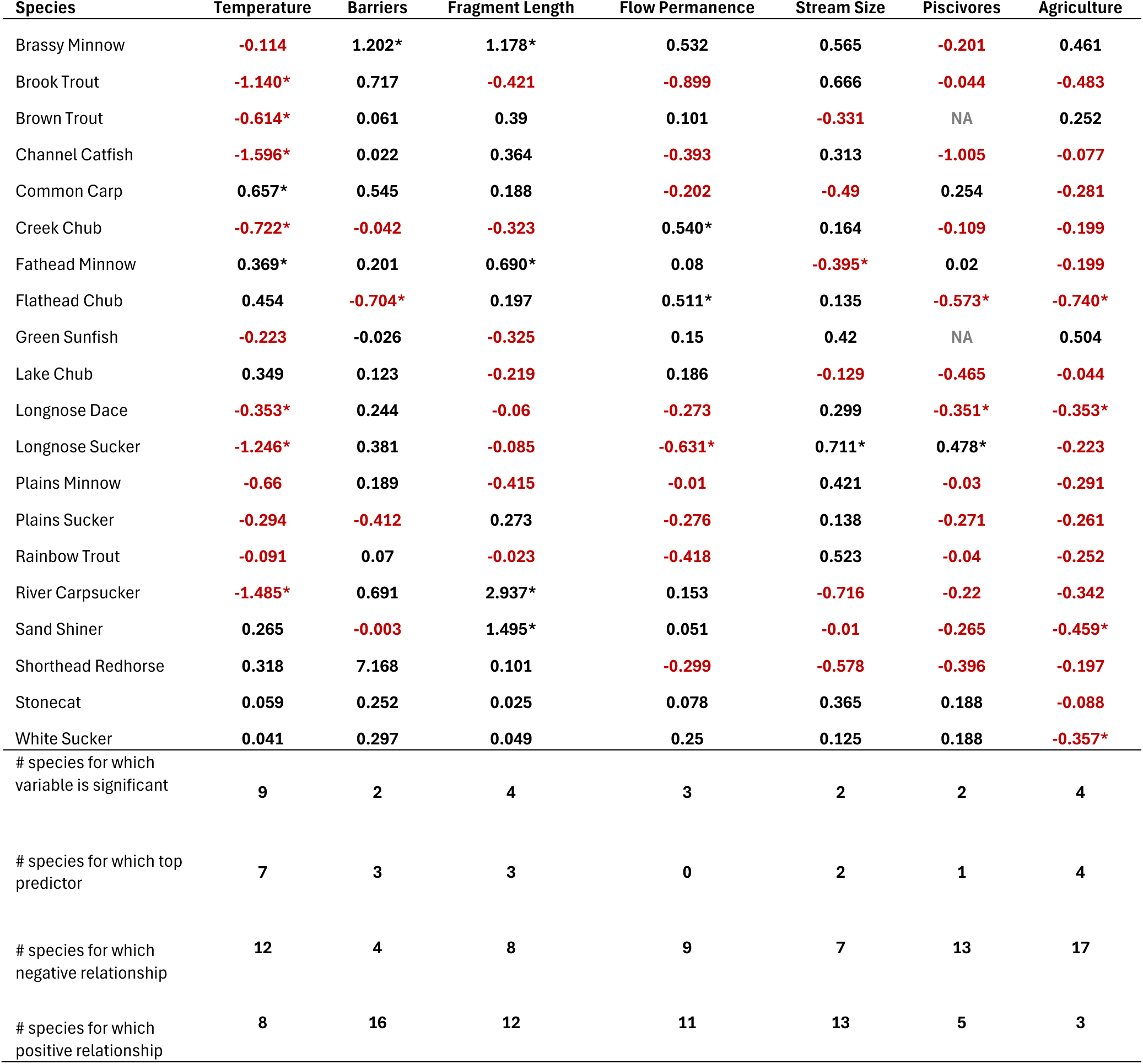
Scaled variable importance scores for individual species’ persistence-colonization models. Models were only run for species found at a minimum of 30 unique sites. Positive relationships between the given variable and the species’ persistence or colonization are shown in black. Negative relationships are shown in red. Brown Trout and Green Sunfish did not include introduced piscivores as a variable due to their inclusion in the predictor itself. * signifies significance at α= 0.10.

Community turnover was composed of approximately equal parts colonization of new species (45%) and loss of historically present species (55%). No individual species was responsible for more than 7% of overall community turnover, but white sucker, fathead minnow, and longnose dace combined accounted for almost 20% (Table S1). Mean August stream temperature was the top predictor of community turnover, native species persistence, and proportional colonization, and agricultural land-use was the second-best predictor (Figure 3A-C). Interactions between temperature and agricultural land-use were also apparent for community turnover and colonization wherein higher levels of agricultural land use led to more turnover and colonization in cooler streams, but not in warmer streams (Figure 3G, I). The same interaction was not apparent (and did not improve model fit) for native species persistence (Figure 3H). Best-fit models for all three community response variables (community turnover, native species persistence, and colonization) included temperature and agricultural land use, and those for native species persistence also included fragment length (Table S2, Figure 3).

**Figure 3.**
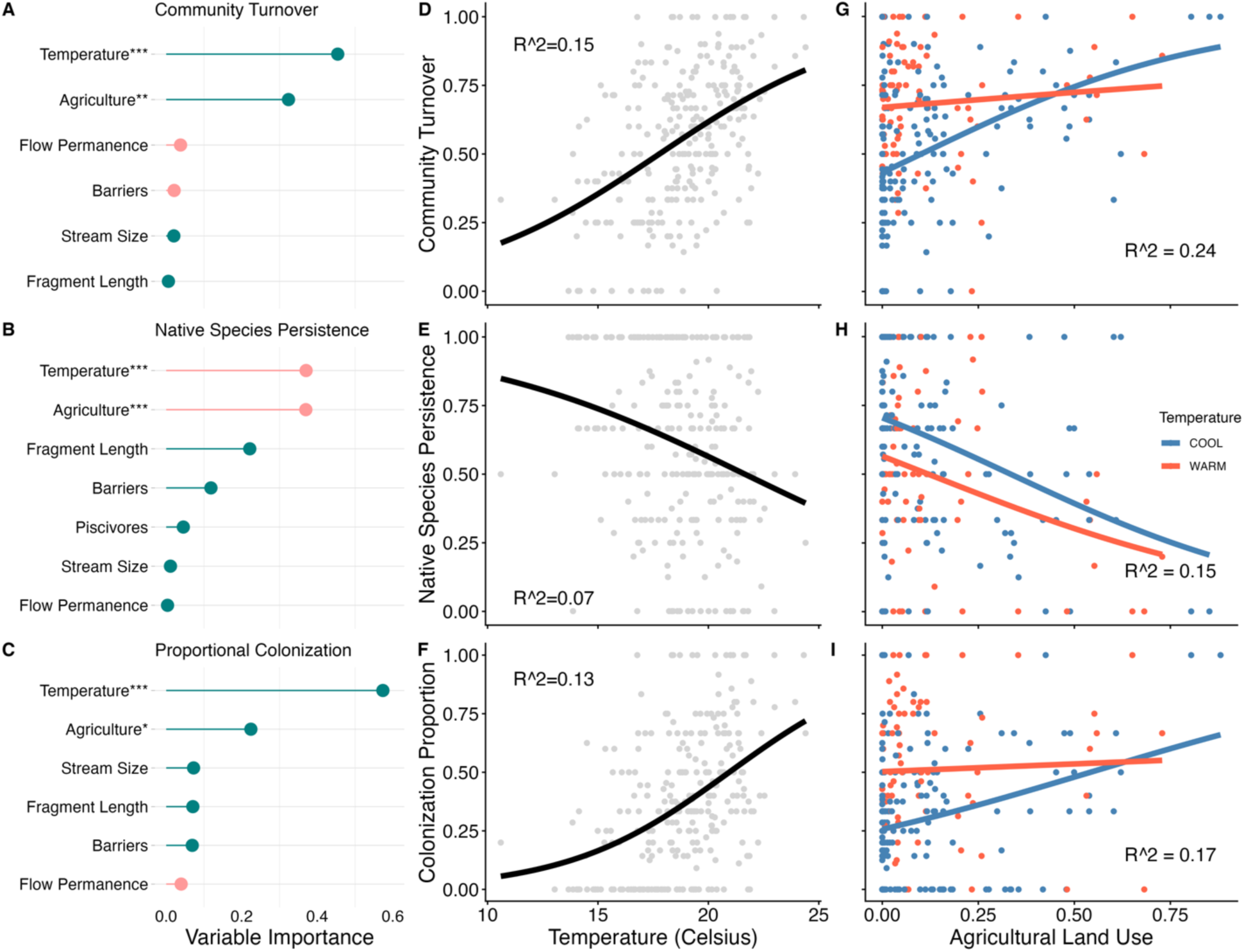
(A-C) Plots of variable importance from generalized linear models for community turnover (A), native species persistence (B), and proportion colonizing species (C). Green indicates a positive correlation with the response variable, and pink indicates a negative correlation. P-values are shown as asterisks (***p <0.01, **p<0.05, *p<0.10). (D-F) Relationships between temperature and the same three response variables. (G-I) Relationships between agricultural land-use and the same three response variables by cool (<20° C) and warm (>20° C) mean August stream temperature.

### Variation in Drivers of Community Change Across Basins

There were significant differences for all three community metrics among basins (Figure 1; ANOVA p-values all less than 0.10). The Upper Missouri River basin consistently exhibited the greatest change (highest mean community turnover and mean colonization and lowest mean native species persistence) followed by the Yellowstone River basin. The Platte River basin consistently exhibited the least amount of community change.

Within major river basins, temperature was the top predictor of community turnover and colonization in the Black Hills, Platte, and Yellowstone River basins while agricultural land use was the top predictor for the Upper Missouri River basin (Figures S1-S3). Top basin-specific predictors for native species persistence were the same for the Black Hills, Platte (both temperature), and Upper Missouri (agricultural land use), but stream fragment length was the top predictor of native species persistence in the Yellowstone River basin.

## DISCUSSION

Community turnover was driven approximately equally by species colonization and species loss. Warm stream temperatures and heavy agricultural land use were the two variables most associated with high community turnover, decreased native species persistence, and increased colonization. There also appears to be an interaction between the two variables such that cool temperatures buffer communities from turnover and colonization by new species, even in areas with moderate agricultural use. In areas with heavy agricultural land-use, community turnover tended to be high, regardless of temperature.

Our results are consistent with the vast literature on the importance of temperature for regulating fish-community structure (e.g., Lyons 1996; Herlihy et al. 2006) and the observed effects of climate warming on fish distributions (Comte et al. 2021a; Kirk and Rahel 2023). Our results are also consistent with studies showing agricultural land use is associated with fish community turnover (Walser and Bart 1999; Chen and Olden 2020). Likely mechanisms underlying this relationship are increased sedimentation, non-point source pollution, floodplain disconnection, stream channelization, and water diversion associated with agricultural land use and negative impacts to fishes (Walser and Bart 1999; EPA 2002). Comte et al. (2021a) demonstrated that climate change and human development interact to reorganize stream fish communities on a global level, and our results corroborate that finding for the northern Great Plains, specifically with stream warming and agricultural land use. As such, conservation of persistent coolwater patches, even in areas with moderate amounts of farming or grazing, may be an important climate adaptation tool.

Importantly, there was spatial variation in the relative importance of these two variables. While temperature was the dominant factor associated with community turnover across most of the study area, agricultural land-use was dominant in the Upper Missouri River basin of northern Montana. This region is heavily impacted by farming, primarily for wheat, and the average proportion of agricultural land use around survey sites was nearly three times higher that of the next most impacted basin (23% in the Upper Missouri vs. 8% in the Yellowstone).

Degraded agricultural lands are often targets for restoration, and interest in the restoration of prairie streams is increasing (Lenhart et al. 2023). Some studies indicate that restoration of habitat or stream connectivity may increase abundances of native prairie fishes by increasing habitat availability (Gido et al. 2023; Clancy et al. *in press*). However, we are unaware of any studies evaluating whether restoration can reverse prairie fish community turnover. As such, protection of existing prairie streams via conservation easement, grazing management, and fencing may the best means of promoting fish community persistence.

### Limited community response to piscivores and streamflow permanence

One of the most surprising findings was the lack of a substantial relationship between community metrics and introduced piscivores and streamflow permanence, variables we expected to be important. For instance, previous studies have indicated that native fishes, especially small, prairie minnows, can be highly susceptible to extirpation by invasive species such as northern pike and smallmouth bass (Stringer 2016; Kirk et al. 2022; Hogberg 2024). However, introduced piscivores did not appear to substantially impact overall native-fish persistence in our study. This supports previous studies that suggest negative community impacts of invasive species may be less important than abiotic conditions at landscape scales, especially in regions with severe environmental conditions such as drought and temperature extremes (Jackson et al. 2001; Booher and Walters 2021; Coulter et al. 2024). We also did not include data on piscivore density which is more biologically meaningful than mere presence-absence when invasive species must reach a threshold abundance before they exert major effects on native species (Quist et al. 2004; Klein et al. 2023). Similarly, piscivores may alter the abundance of prey species and thus alter community composition without causing extirpations. In addition, we only examined whether introduced piscivores were present in the contemporary time period - it is possible some of the effects of these species on native species had already occurred during the historical time period. Despite these potential shortcomings, the relatively low importance of piscivores for community metrics aligns with previous studies indicating that abiotic conditions act as a higher-level, biogeographic “filter” for communities (*sensu* Poff 1997; Quist et al. 2005). In the northern Great Plains, we expect that naturally high levels of intermittency, thermal variability, and turbidity limits the spread and abundance of piscivores, thus limiting their influence (e.g., Hogberg 2024; Coulter et al. 2024).

Streamflow permanence was also surprisingly unexplanatory for community change metrics. Stream drying is known to cause recruitment failure and extirpations of native prairie fishes in the central and southern Great Plains (Falke et al. 2010; Perkin et al. 2015; Perkin et al. 2017). It is possible prairie stream fish communities that are already adapted to extreme cycles of flooding and drying, are largely able to tolerate the current shifts in streamflow timing and magnitude. Alternatively, our metric of streamflow permanence may not accurately characterize streamflow permanence changes in areas heavily impacted by agriculture because it comes from a model that did not include water-diversion predictors or change in streamflow magnitude or timing (Sando et al. 2022).

### Individual species show highly variable responses to environmental factors

Top predictors for individual species’ persistence and colonization were highly variable. While temperature and agriculture were top predictors for some species, they were not as uniformly important as with community metrics. The nine declining native species that we modelled had particularly variable top predictors of persistence and turnover that included temperature (longnose dace, longnose sucker, plains minnow); barriers (brassy minnow, plains sucker); fragment length (fathead minnow); piscivores (flathead chub, lake chub); and agricultural land use (longnose dace, white sucker). Species also showed varying relationships to aspects of stream connectivity (presence of a downstream barrier, stream fragment length, and flow permanence) with some species benefiting from fragmentation and some declining. For example, both brassy minnow and fathead minnow were positively associated with the presence of a downstream barrier and long fragments, likely indicating that downstream barriers protect the species from competition or predation by colonizing species. Flathead chub and plains sucker showed the opposite relationship and were negatively associated with downstream barriers. Flathead chub is a pelagic broadcast spawner, and the negative relationship of such fishes with in-stream barriers is well established (Perkin and Gido 2011; Perkin et al. 2015). These opposing relationships demonstrate that aggregated community metrics can obscure species-specific relationships.

### Implications for Climate Change Refugia

Individual species trends in our study generally align with future climate-change predictions from regional studies (Table S3). This demonstrates that climate-induced range shifts for many species are well under way, and points to the need for delineating and conserving climate refugia for native fish assemblages. The highly variable drivers of individual species change, especially for declining fishes, indicates that studies of climate vulnerability and delineation of climate change refugia will be most accurate when completed for individual species. However, a generalization that emerged from our study is that warm stream temperatures and high proportions of agricultural land-use were associated with high community turnover, low native species persistence, and high rates of colonization. Thus, watersheds with cool water and low agricultural land-use will likely retain a majority of native, prairie stream fishes and would be good climate refugia.

Our results also add important context to the connectivity vs. coldwater refugia debate (*sensu* Armstrong et al. 2019; Isaak and Young 2023). Overall, our results suggest cooler summer temperatures are the most important variable for the persistence of native prairie fishes and lower rates of species colonization. However, for some species, stream connectivity (i.e., no barriers or long stream fragment lengths) is the most important predictor of persistence. From a fish community perspective, a management dichotomy prioritizing either cool temperatures or stream connectivity is not productive. Because different species require different temperatures and levels of connectivity or isolation from introduced species (including natural isolation due to stream intermittency), basins prioritized for conservation will benefit from containing a diverse portfolio of water temperatures, unfragmented streams, perennial and intermittent habitats, and isolated reaches. The useful question is not whether cold temperatures or connectivity should be prioritized, but what amount of coldwater habitat and isolation is needed to support healthy fish communities?

### Management Implications

Long-term community persistence in our study was associated with cool water temperatures and low levels of agricultural impact. Our results highlight the potential for protection (and possibly restoration) of coolwater habitats to buffer fish communities from the impacts of low to moderate agricultural land-use. Specific management actions include protecting instream flow rights, late summer hypolimnetic dam releases, riparian protection and restoration, and mapping and protection of coolwater refugia. However, individual species have highly variable relationships to the environmental factors we examined, so conservation plans for at-risk species could benefit from species-specific environmental requirements.

## ACKNOWLEDGEMENTS

We first thank the many state, federal and university biologists who collected data included in our study. We also thank Elizabeth Rieger, Toby Covill, Chris Taylor, Ella Humphrey, and Nolan Weatherby for their dedicated help in the field. Conversations with Jeff Baldock helped refine the context of this study. Yoichiro Kanno, Di Yang, and Fabian Nippgen provided helpful comments that improved the manuscript. Funding was provided by the North Central Climate Adaptation Science Center and Wyoming Landscape Conservation Initiative. Field collections were made under the conditions of UW IACUC protocol 20211001AW00518-02, and collector’s permits for Wyoming (#1388), Montana (27-2022, 38-2023, and 09-2024), and North Dakota (#19905). Any use of trade, firm or product names is for descriptive purposes only and does not imply endorsement by the U.S. Government.

## Data availability

the dataset used in this study can be found as USGS data release (www.doi.org/10.5066/P1YXQRBD) and at https://doi.org/10.5281/zenodo.14783076.

Associated R code can be found at https://github.com/niallgclancy/persistence

---

**Figure S1.**
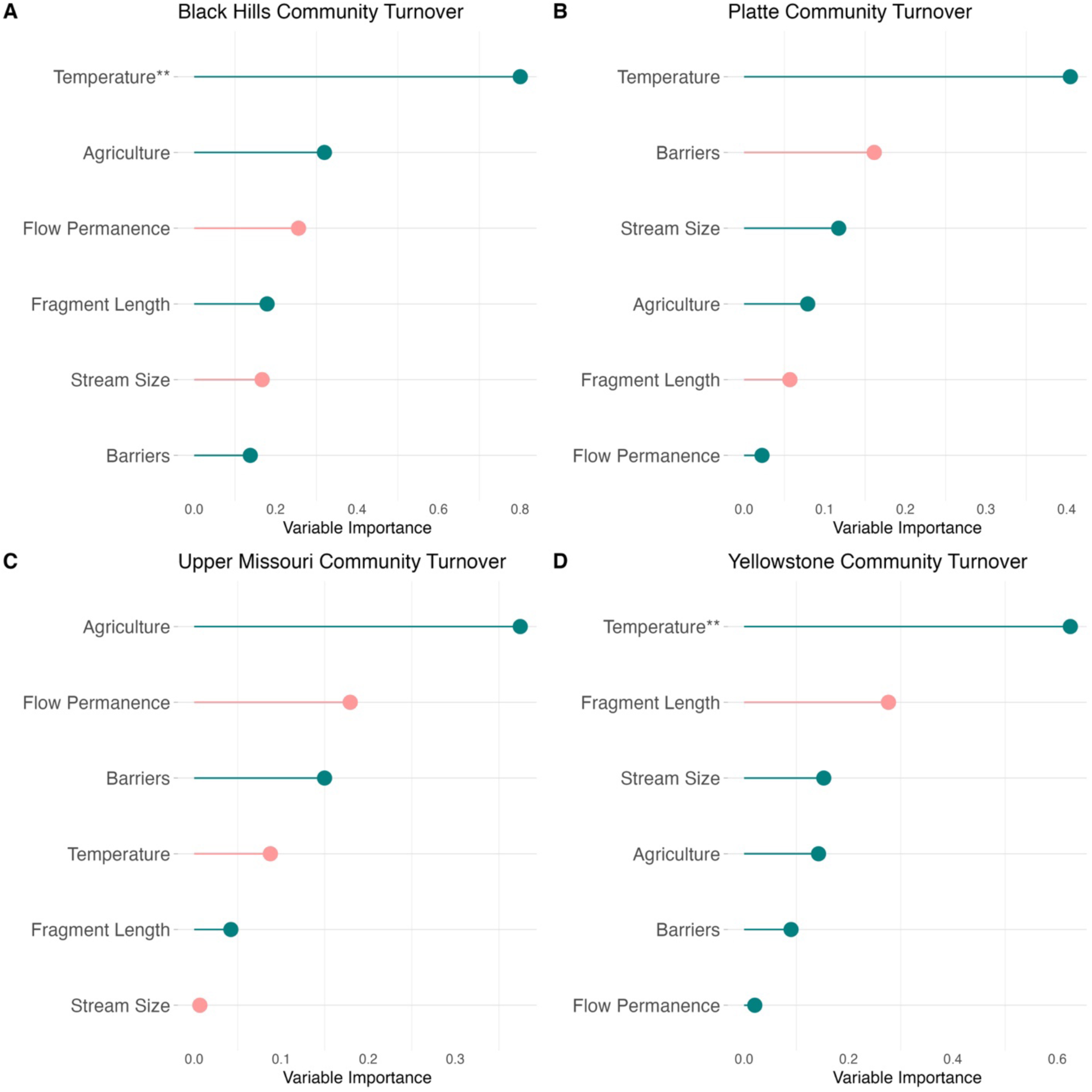
Plots of variable importance from generalized linear models for community turnover by major drainage basin. Green indicates a positive correlation with the response variable, and pink indicates a negative correlation. P-values are shown as asterisks (***p <0.01, **p<0.05, *p<0.10).

**Figure S2.**
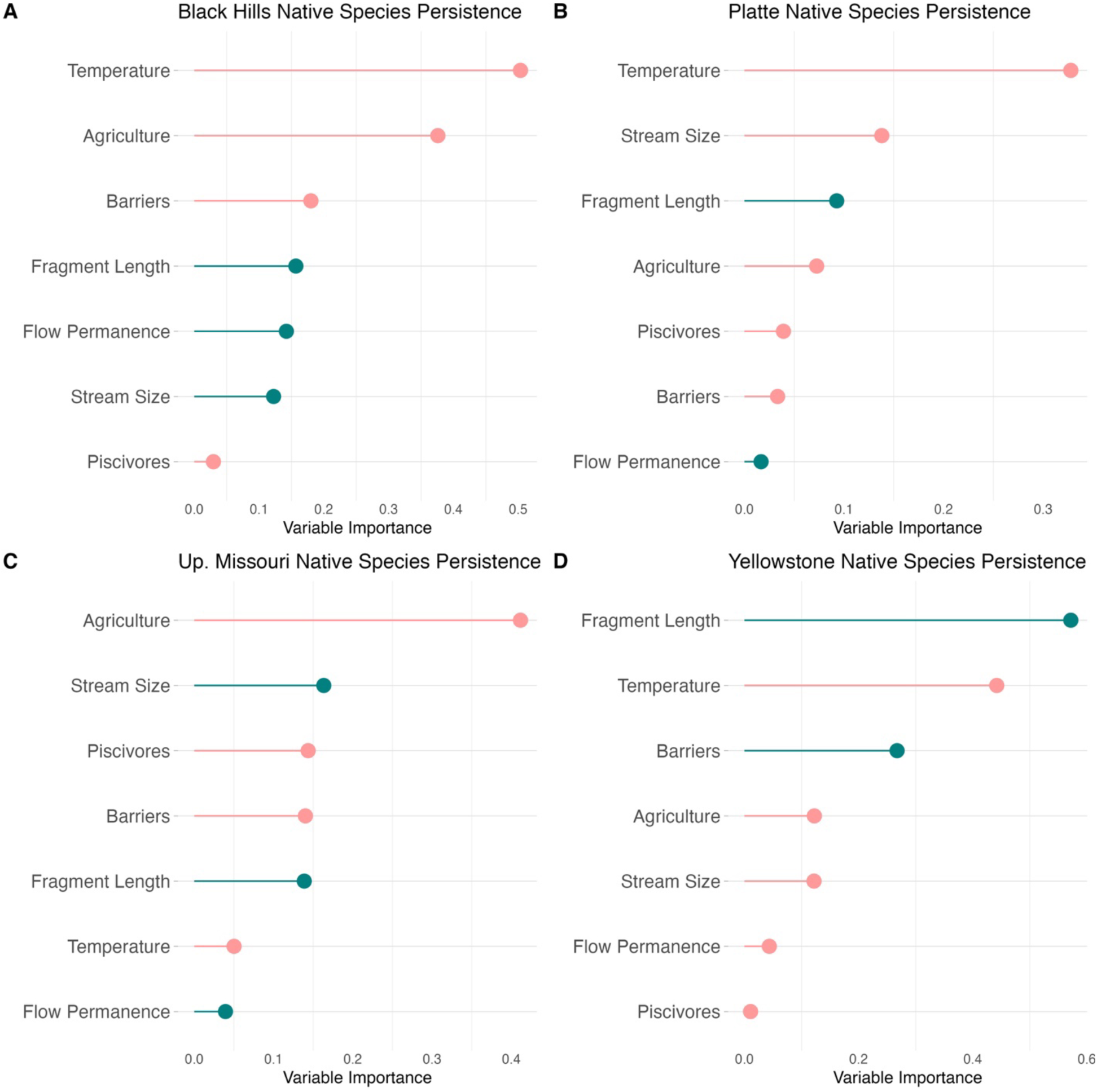
Plots of variable importance from generalized linear models for native species persistence by major drainage basin. Green indicates a positive correlation with the response variable, and pink indicates a negative correlation. No covariate p-values were below 0.10.

**Figure S3.**
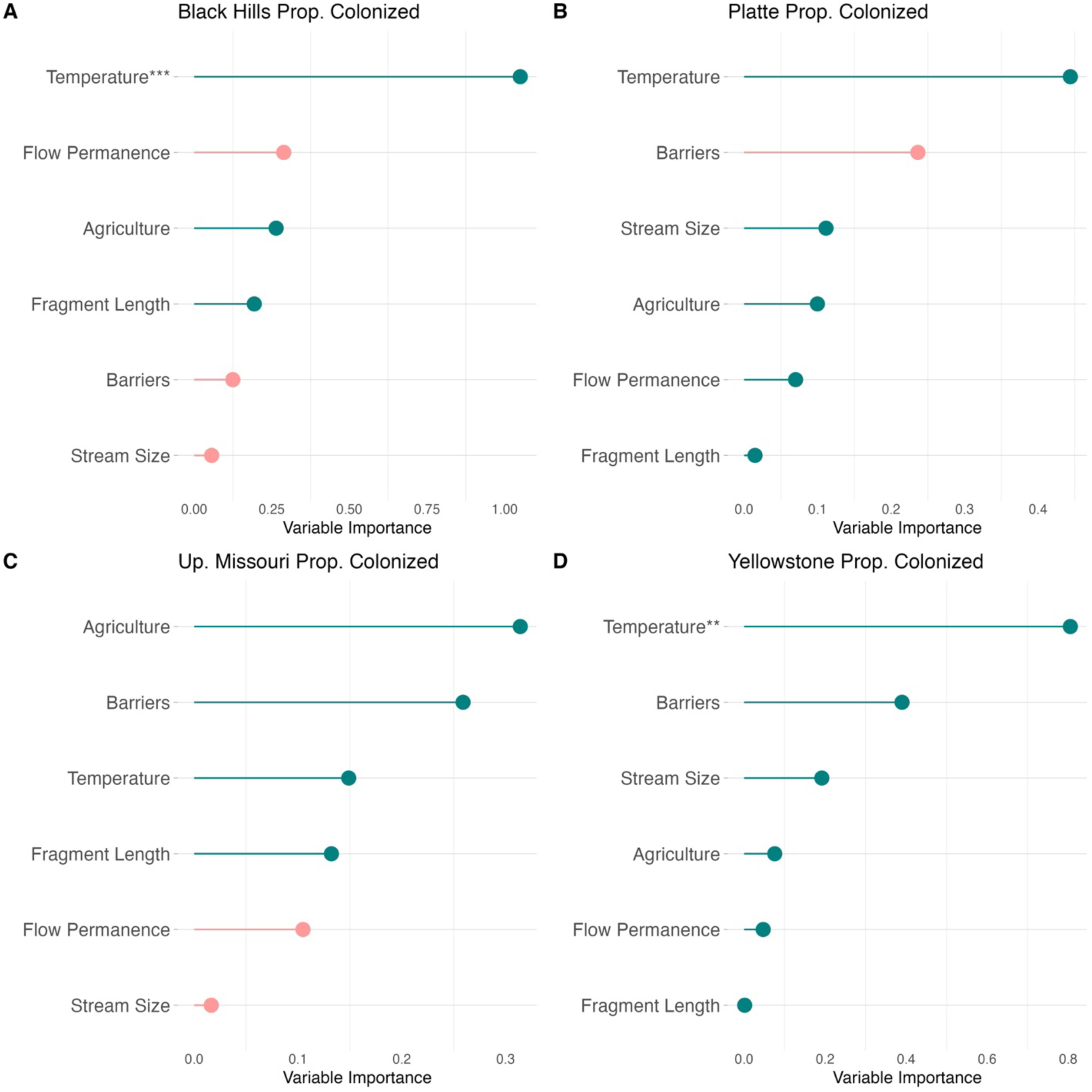
Plots of variable importance from generalized linear models for proportional colonization by major drainage basin. Green indicates a positive correlation with the response variable, and pink indicates a negative correlation. P-values are shown as asterisks (***p <0.01, **p<0.05, *p<0.10).

**Table S1.** Proportion of mean community turnover accounted for by colonization or loss of individual species.

| Species | Mean Proportion of Turnover<br>Due to Individual Species |
| --- | --- |
| White Sucker | 0.067 |
| Fathead Minnow | 0.066 |
| Longnose Dace | 0.062 |
| Longnose Sucker | 0.042 |
| Green Sunfish | 0.040 |
| Sand Shiner | 0.037 |
| Common Carp | 0.037 |
| Creek Chub | 0.034 |
| Stonecat | 0.032 |
| Brassy Minnow | 0.030 |
| Plains Minnow | 0.029 |
| Black Bullhead | 0.029 |
| Plains Sucker | 0.027 |
| Shorthead Redhorse | 0.027 |
| Lake Chub | 0.026 |
| Flathead Chub | 0.026 |
| Channel Catfish | 0.026 |
| Rainbow Trout | 0.025 |
| River Carpsucker | 0.025 |
| Brook Stickleback | 0.023 |
| Brown Trout | 0.022 |
| Northern Plains Killifish | 0.021 |
| Iowa Darter | 0.017 |
| Brook Trout | 0.016 |
| Smallmouth Bass | 0.015 |
| Goldeye | 0.014 |
| Western Silvery Minnow | 0.012 |
| Johnny Darter | 0.012 |
| Northern Pike | 0.012 |
| Northern Redbelly Dace | 0.011 |
| Largemouth Bass | 0.009 |
| Bigmouth Shiner | 0.009 |
| Plains Topminnow | 0.009 |
| Common Shiner | 0.009 |
| Yellow Perch | 0.009 |
| Hornyhead Chub | 0.008 |
| Hybrid *Oncorhynchus* | 0.006 |
| Rock Bass | 0.006 |
| Red Shiner | 0.006 |
| Central Stoneroller | 0.006 |
| Golden Shiner | 0.005 |
| Walleye | 0.005 |

**Table S2.** The top five (lowest AIC) models for each of the three community change metrics. * indicates that an interaction between the two terms is included in the model, and + indicates no interaction is included.

| Model Covariates | Mean Squared Error (MSE) | Root Mean Square Error (RMSE) | AIC | Δ MSE | Δ AIC |
| --- | --- | --- | --- | --- | --- |
| **Community Turnover** |  |  |  |  |  |
| TEMP * AGRI | 0.0477 | 0.218 | 334.8 | 0.0000 | 0.0000 |
| TEMP * PERM * AGRI | 0.0482 | 0.220 | 341.3 | 0.0006 | 6.4576 |
| TEMP * LENGTH * AGRI | 0.0483 | 0.220 | 342.4 | 0.0006 | 7.5730 |
| TEMP * DAM * AGRI | 0.0485 | 0.220 | 342.4 | 0.0008 | 7.5301 |
| TEMP * SIZE * AGRI | 0.0488 | 0.221 | 342.1 | 0.0012 | 7.2758 |
| **Native Species Persistence** |  |  |  |  |  |
| TEMP + LENGTH + AGRI | 0.085 | 0.292 | 350.5 | 0.0001 | 0.0000 |
| TEMP + DAM + LENGTH + AGRI | 0.085 | 0.292 | 351.0 | 0.0000 | 0.4979 |
| TEMP * DAM * LENGTH * PERM * AGRI | 0.085 | 0.292 | 372.1 | 0.0001 | 21.5820 |
| TEMP + LENGTH + SIZE + AGRI | 0.086 | 0.293 | 352.0 | 0.0005 | 1.5089 |
| TEMP + DAM + LENGTH + SIZE + AGRI | 0.086 | 0.293 | 352.9 | 0.0005 | 2.3837 |
| TEMP + DAM + LENGTH + PERM + AGRI | 0.086 | 0.293 | 352.9 | 0.0005 | 2.3959 |
| **Colonization** |  |  |  |  |  |
| TEMP * AGRI | 0.067 | 0.258 | 306.4 | 0.0000 | 0.0000 |
| TEMP * LENGTH * AGRI | 0.068 | 0.260 | 314.6 | 0.0010 | 8.2461 |
| TEMP + SIZE + AGRI | 0.068 | 0.261 | 307.8 | 0.0013 | 1.4214 |
| TEMP + AGRI | 0.068 | 0.261 | 306.7 | 0.0013 | 0.3659 |
| TEMP + LENGTH + AGRI | 0.068 | 0.261 | 308.3 | 0.0014 | 1.9477 |

**Table S3.**
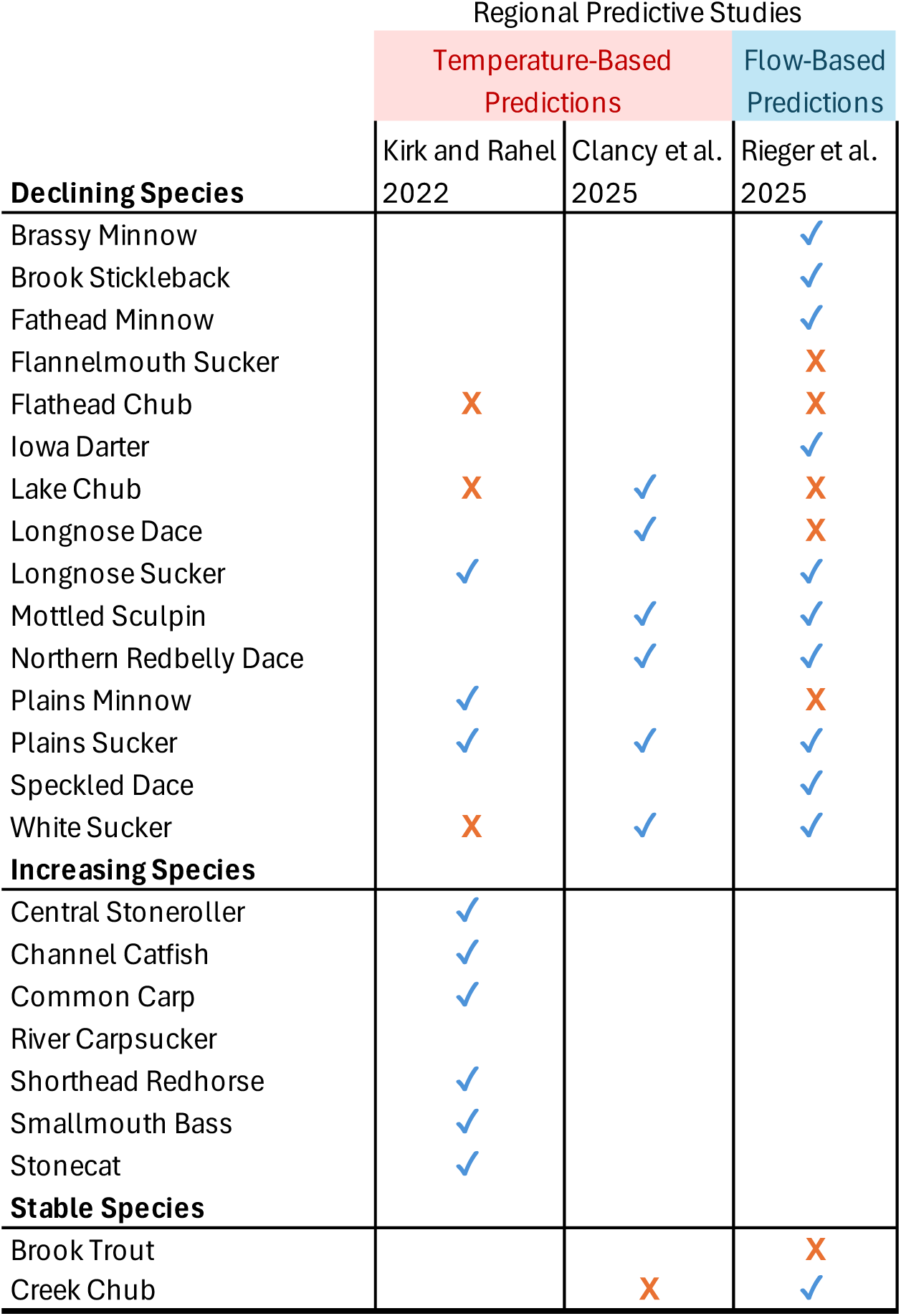
Comparison of species’ observed trends with three studies that predicted species’ responses to climate change in the same region. Blue checks indicate observed trends corroborate predictions and orange Xs indicate observed trends differed from predictions. A blank indicates the study did not include that species. Clancy et al. (2025) and Rieger et al. (2026) only evaluated potential declines (not colonization probabilities), so we only compared species predicted to substantially decline (20% or more) by those studies with observed trends.

